# A preparatory cranial potential for saccadic eye movements in macaque monkeys

**DOI:** 10.1101/2025.01.16.633363

**Authors:** Steven P. Errington, Jeffrey D. Schall

## Abstract

Response preparation is accomplished by gradual accumulation in neural activity until a threshold is reached. In humans, such a preparatory signal, referred to as the lateralized readiness potential, can be observed in the EEG over sensorimotor cortical areas before execution of a voluntary movement. Although well-described for manual movements, less is known about preparatory EEG potentials for saccadic eye movements in humans and nonhuman primates. Hence, we describe a lateralized readiness potential over the frontolateral cortex in macaque monkeys. Homologous to humans, we observed lateralized electrical potentials ramping before the execution of both rewarded and non-rewarded contralateral saccades. This potential parallels the neural spiking of saccadic movement neurons in the frontal eye field, suggesting that it may offer a non-invasive correlate of intracortical spiking activity. However, unlike neural spiking in the frontal eye field, polarization in frontolateral channels did not distinguish between saccade generation and inhibition. These findings provide new insights into non-invasive electrophysiological signatures of saccadic preparation in nonhuman primates, highlighting the potential of EEG measures to bridge invasive neural recordings and non-invasive studies of eye movement control in humans.

## INTRODUCTION

Intentional control over our actions emerges from preparatory processes that ensures coordinated and purposeful interactions with our surroundings. This preparatory activity manifests neurally through a slow buildup in activity, which accumulates over time (Hanes & Schall, 1996; Riehle & Requin, 1989; Tanji & Evarts, 1976; Weinrich et al., 1984). In humans, this process has been well described through the lateralized readiness potential (de Jong et al., 1988; Gratton et al., 1988; Smulders et al., 2012). The lateralized readiness potential is a gradual buildup of electrical polarization observed in EEG signals over the motor cortex. This potential occurs one or two hundred milliseconds before a voluntary movement is executed, and thought to reflect the preparatory processes in the motor cortex for generating an action (Deecke et al., 1976).

Although the lateralized readiness potential is usually described in the context of a voluntary body movement, such as an arm or leg, similar lateralized EEG components have been observed during the preparation of saccadic eye movements (Balaban & Weinstein, 1985; Brickett et al., 1984; Evdokimidis et al., 1991; Evdokimidis et al., 1992; Everling, Krappmann, & Flohr, 1997; Everling, Krappmann, Spantekow, et al., 1997; Kurtzberg & Vaughan, 1982; Moster & Goldberg, 1990; Thickbroom & Mastaglia, 1985). Interestingly, these non-invasive event-related potentials show similar dynamics to spiking activity observed in invasive neurophysiological studies of the the macaque frontal eye field (FEF) - a region in the frontal cortex of the brain that plays a crucial role in the preparation and execution of saccadic eye movements (Bruce & Goldberg, 1985; Hanes et al., 1998). FEF neurons show a specific type of ramping activity known as “ramp-to-threshold” which is interrupted when saccades are inhibited (Hanes et al., 1998; Hanes & Schall, 1996). This movement-related activity occurs prior to both externally cued saccades (Bruce & Goldberg, 1985) and spontaenous, voluntary saccades (Sendhilnathan et al., 2021, but see Bizzi, 1968).

Previous work has demonstrated that monkeys have homologues of human cognitive ERP components, such as: N2pc (Cohen et al., 2009; Heitz et al., 2010; Purcell et al., 2013; Woodman et al., 2007),contralateral delay activity (Reinhart et al., 2012), ERN/Pe (Godlove et al., 2011; Sajad et al., 2019), and N2/P3 (Sajad et al., 2022). These observations have helped establish animal models that can be used to investigate the neural mechanisms underlying these event-related potentials (Herrera et al., 2023; Herrera et al., 2024; Herrera et al., 2022). Here, we expand on this literature to describe a pre-saccadic lateralized readiness-potential in EEG signals recorded over frontolateral cortex.

## MATERIALS & METHODS

### Experimental models and subject details

Data was collected from one male bonnet macaque (Eu, Macaca Radiata, 8.8 kg) and one female rhesus macaque (X, Macaca Mulatta, 6.0 kg) performing a countermanding task (Godlove et al., 2014; Hanes & Schall, 1995). All procedures were approved by the Vanderbilt Institutional Animal Care and Use Committee in accordance with the United States Department of Agriculture and Public Health Service Policy on Humane Care and Use of Laboratory Animals.

### Animal care and surgical procedures

Surgical details have been described previously (Godlove et al., 2011). Briefly, magnetic resonance images (MRIs) were acquired with a Philips Intera Achieva 3T scanner using SENSE Flex-S surface coils placed above or below the animal’s head. T1-weighted gradient-echo structural images were obtained with a 3D turbo field echo anatomical sequence (TR = 8.729 ms; 130 slices, 0.70 mm thickness). These images were used to ensure that Cilux recording chambers were placed in the correct area (Crist Instruments). Chambers were implanted normal to the cortex (Monkey Eu: 17o; Monkey X: 9o; relative to stereotaxic vertical) centered on the mid-line, 30 mm (Monkey Eu) and 28mm (Monkey X) anterior to the interaural line.

### EEG processing and data acquisition

Electrophysiology data were processed with unity-gain high-input impedance head stages (HST/32o25-36P-TR, Plexon). All data were streamed to a single data acquisition system (MAP, Plexon, Dallas, TX). Time stamps of trial events were recorded at 500 Hz. Eye position data were streamed to the Plexon computer at 1 kHz using an EyeLink 1000 infrared eye-tracking system (SR Research, Kanata, Ontario, Canada).

### Saccade stop-signal task

The saccade stop-signal (countermanding) task utilized in this study has been widely used previously (Cabel et al., 2000; Colonius et al., 2001; Godlove & Schall, 2016; Hanes & Carpenter, 1999; Hanes & Schall, 1995; Kornylo et al., 2003; Morein-Zamir & Kingstone, 2006; Thakkar et al., 2011; Thakkar et al., 2015; Verbruggen et al., 2019; Walton & Gandhi, 2006; Wattiez et al., 2016). Briefly, trials were initiated when monkeys fixated on a central point. Following a variable period, the center of the fixation point was removed leaving an outline. At this point, a peripheral target was presented simultaneously on either the left or right hand of the screen. In this study, one target location was associated with a larger magnitude of fluid reward. The lower magnitude reward ranged from 0 to 50% of the higher magnitude reward amount. This proportion was adjusted to encourage the monkey to continue responding to both targets. The stimulus-response mapping of location-to-high reward changed across blocks of trials. Block length was adjusted to maintain performance at both targets, with the number of trials in each block determined by the number of correct trials performed. In most sessions, the block length was set at 10 to 30 correct trials. Erroneous responses led to repetitions of a target location, ensuring that monkeys did not neglect low-reward targets in favor of high-reward targets – a phenomenon demonstrated in previous implementations of asymmetrically rewarded tasks (Kawagoe et al., 1998).

In most of the trials, the monkey was required to make an eye movement toward this target (no-stop trials). However, in a proportion of trials the center of the fixation point was re-illuminated (stop-signal trials); this stop-signal appeared at a variable time after the target had appeared (stop-signal delay; SSDs). An initial set of SSDs, separated by either 40 or 60 ms, were selected for each recording session. The delay was then manipulated through an adaptive staircasing procedure in which stopping difficulty was based on performance. When a subject failed to inhibit a response, the SSD was decreased by a random step to increase the likelihood of success on the next stop trial. Similarly, when subjects were successful in their inhibition, the SSD was increased to reduce the likelihood of success on the next stop trial. This procedure was employed to ensure that subjects failed to inhibit action on approximately 50% of all stop-signal trials. On no-stop trials, the monkey was rewarded for making a saccade to the target. In stop-signal trials, the monkey was rewarded for withholding the saccade and maintaining fixation on the fixation spot. Following a correct response, an auditory tone was sounded 600ms later, and followed by a high or low fluid reward, depending on the stimulus-response mapping.

### Data collection protocol

An identical daily recording protocol across monkeys and sessions was carried out. In each session, the monkey sat in an enclosed primate chair with their head restrained 45cm from a CRT monitor (Dell P1130, background luminance of 0.10 cd/m2). The monitor had a refresh rate of 70Hz, and the screen subtended 46 deg x 36 deg of the visual angle. Eye position data were collected at 1 kHz using an EyeLink 1000 infrared eye-tracking system (SR Research, Kanata, Ontario, Canada). This was streamed to a single data acquisition system (MAP, Plexon, Dallas, TX) and amalgamated with other behavioral and neurophysiological data. After advancing the electrode array to the desired depth, they were left for 3 to 4 hours until recordings stabilized across contacts. This led to consistently stable recordings. Once these recordings stabilized, an hour of resting-state activity in near-total darkness was recorded. This was followed by the passive presentation of visual flashes followed by periods of total darkness in alternating blocks. Finally, the monkey then performed approximately 2000 to 3000 trials of the saccade countermanding (stop-signal) task.

### Bayesian modeling of stop-signal performance

As performance on the stop-signal task can be considered as the outcome of a race between a GO and STOP process, then a stop-signal reaction time (SSRT) can be calculated (Logan & Cowan, 1984). This value can be considered as the latency of the inhibitory process that interrupts movement preparation. Stop-signal reaction time was estimated using a Bayesian parametric approach (Matzke, Dolan, et al., 2013; Matzke, Love, et al., 2013). Compared to classical methods of calculating SSRT (integration-weighted method, Logan and Cowan (1984)), this approach allows for a distribution of SSRT to be derived by using the distribution of reaction times on no-stop trials, and by considering reaction times on non-canceled trials as a censored no-stop response time (RT) distribution. Furthermore, this model also allows for the estimation of the probability of trigger failures for a given session (Matzke et al., 2017). Individual parameters were estimated for each session. The priors were bounded uniform distributions (μ_Go_, μ_Stop_: U (0.001, 1000); σ_Go_, σ_Stop_: U (1, 500) τ_Go_, τ_Stop_: U (1, 500); p_TF_: U (0,1)). The posterior distributions were estimated using Metropolis-within-Gibbs sampling and we ran multiple (3) chains. We ran the model for 5000 samples with a thinning of 5.

## RESULTS

### Monkeys produced planned saccades to targets and inhibited them when instructed

We acquired 33,816 trials across 29 sessions from two macaques (Eu: 11,583; X: 22,233) performing the saccade stop-signal (countermanding) task (Fig. 1A). In the majority of these trials (∼60%), monkeys were required to make a saccade to a target on the left- or right-hand side of the screen. Across all sessions, the median RT to left (m = 267.7 ± 6.8 ms) targets was significantly faster than that to right (m = 248.0 ± 5.8 ms) targets (Mann-Whitney U Test: z = 2.131, p = 0.033). To account for these differences, we only included trials in which the RT was in a mutual range for both directions.

**Fig. 1.**
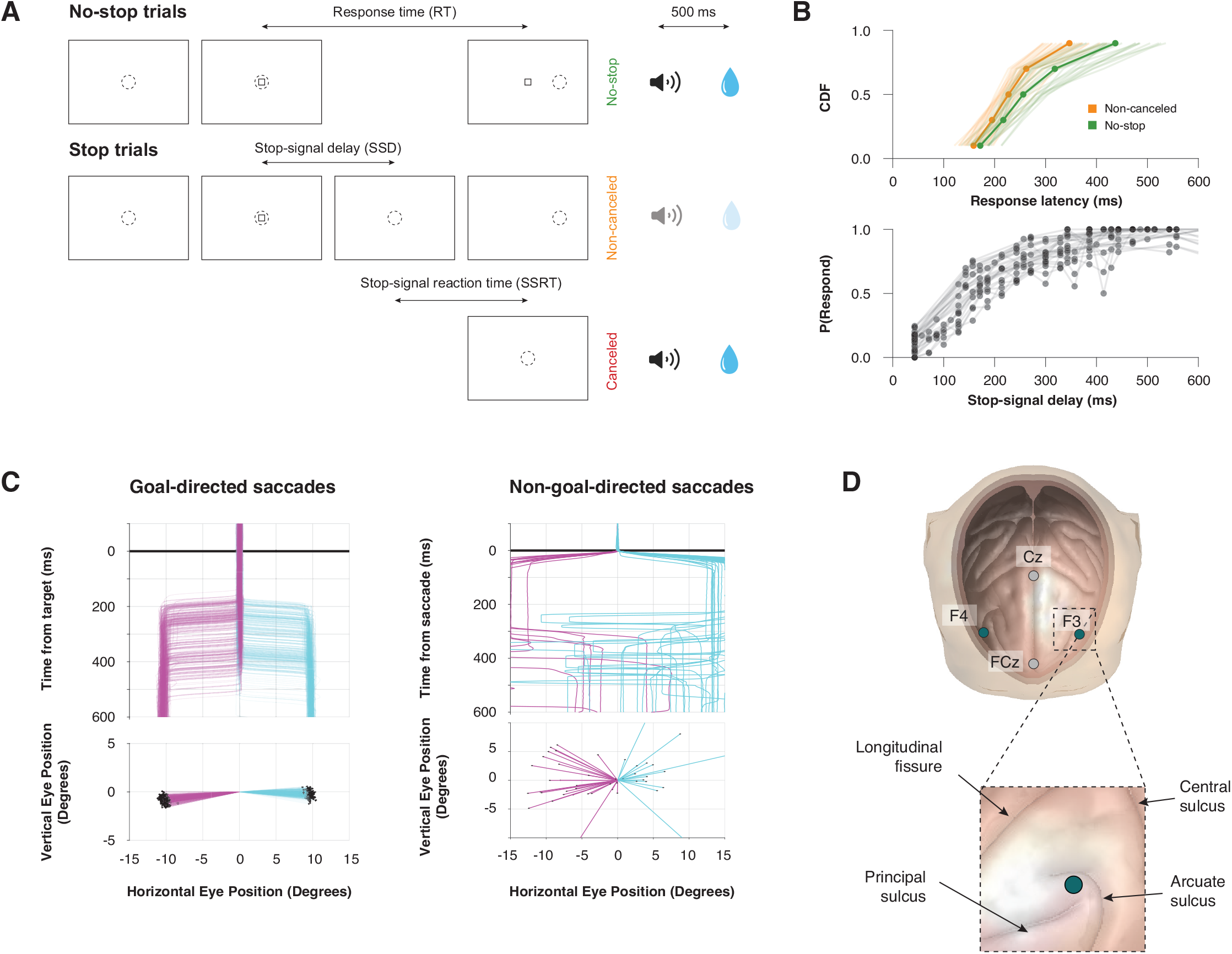
Experimental procedures. **A**. Saccade-countermanding task. Monkeys initiated trials by fixating on a central point. After a variable time, the center of the fixation point was extinguished. A peripheral target was presented simultaneously at one of two possible locations. On no-stop-signal trials monkeys were required to shift gaze to the target, whereupon after 600 ± 0 ms a high-pitched auditory feedback tone was delivered, and 600 ms later fluid reward was provided. On stop-signal trials (∼40% of trials), after the target appeared the center of the fixation point was re-illuminated after a variable stop-signal delay, which instructed the monkey to cancel the saccade in which case the same high-pitched tone was presented after a 1,500 ± 0 ms hold time followed, after 600 ± 0 ms by fluid reward. Stop-signal delay was adjusted such that monkeys successfully canceled the saccade in ∼50% of trials. In the remaining trials, monkeys made non-canceled errors which were followed after 600 ± 0 ms by a low-pitched tone, and no reward was delivered. Monkeys could not initiate trials earlier after errors. **B**. Countermanding behavior. Top: cumulative distribution function of response latencies on no-stop (green) and non-canceled (yellow) trials. Response latencies on non-canceled trials were faster than those on no-stop trials. Bottom: inhibition function plotting the probability of responding across stop-signal delays. **C**. Goal and non-goal directed saccades. Monkeys made both leftward (magenta) and rightward (cyan) saccades with similar properties, in response to the target (goal-directed, left) and spontaneously (non-goal-directed, right). Examples are plotted from one representative example session. Goal-directed sac-cades are shown relative to a central fixation spot and aligned on target onset. Non-goal-directed saccades are shown relative to the pre-saccade eye position, aligned on the time a saccadic eye movement was detected within the intertrial interval. **D**. EEG electrode position. Monkeys were fitted with four EEG electrodes located over the central sulcus (Cz), the medial frontal cortex (FCz), and the frontolateral cortex (F3 & F4). The position of the F3/F4 electrode is estimated to lie over the frontal eye field.

In a small proportion of trials, the appearance of a visual stop-signal instructed the monkey to inhibit the planned saccade. Both monkeys exhibited typical sensitivity to the stop-signal: firstly, response latencies on non-canceled (error) trials were faster than those on no-stop trials (Fig. 1B, top); secondly, the probability of failing to cancel and executing an erroneous saccade was greater at longer stop-signal delays (Fig. 1B, bottom). These two observations validated the assumptions of the independent race model (Logan & Cowan, 1984), allowing us to estimate the stop-signal reaction time (SSRT), the time needed to cancel to partially prepared saccade.

In addition to rewarded, goal-directed saccades (Fig. 1C, left), monkeys also generated spontaneous saccades (Fig. 1C, right) during the inter-trial interval (ITI), when no visual stimuli were displayed, and no rewards could be earned. Here, we limit our analyses to saccades in this period that shared similar kinematic characteristics to the goal-directed saccades: namely, we included saccades that were directed towards the left (180 ± 45°) or right (0 ± 45°) from their point of initiation, which exceeded an amplitude of 15°. Although non-goal directed saccades exhibited higher variability in their characteristics, both types of saccades displayed notable similarities, allowing for comparison of neural signals between the two types.

### Event related potentials over lateral frontal cortex ramp until saccade execution

Event-related potentials (ERPs) were recorded with frontolateral electrodes, homologous to F3 and F4 in the human 10-20 system, during the stop-signal task (Fig. 1D). These contacts were located over the dorsal limb of the arcuate sulcus. Like the lateralized readiness potential reported in humans prior to a voluntary manual movement, we observed a progressive polarization of the EEG in trials in which a goal-direct saccade towards a target was correctly generated (Fig. 2A). This preparatory saccade potential shows a clear potential shift beginning approximately 300 ms prior to saccade onset, which reaches peak amplitude at the time of saccade execution.

**Fig. 2.**
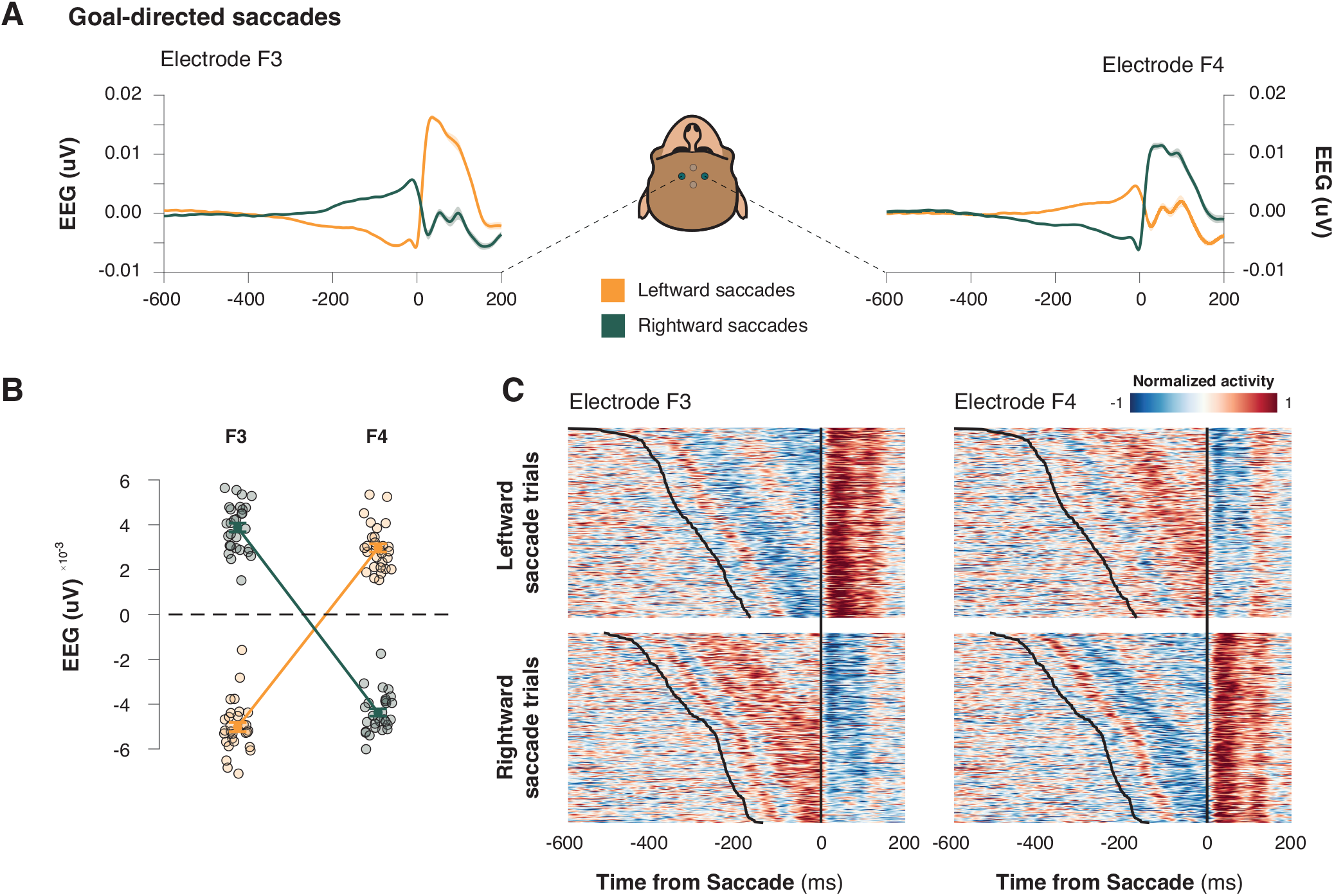
A lateralized readiness potential for goal-directed saccades. **A**. Trial-averaged event-related potentials from an example session for left (F3) and right (F4) frontolateral electrodes during leftward (orange) and rightward (green) goal-directed saccades. **B**. Scatter plot illustrating mean EEG amplitudes prior to the saccade at left (F3) and right (F4) frontolateral electrode sites, highlighting lateralization in saccadic activity. **C**. Heatmaps of normalized activity aligned within an example, representative session. Activity is aligned to saccade onset for left (F3) and right (F4) frontolateral electrodes, for leftward (top) and rightward (bottom) saccades.

Figure 2A displays the grand average ERP waveforms for contralateral and ipsilateral saccades, plotted separately for electrode F3 and F4, locked on the time of saccade. A positively accumulating potential was observed prior to the execution of a saccade in the contralateral direction and a negatively accumulating potential was found prior to the execution of a saccade in the ipsilateral direction (Fig. 2B, Repeated-measures ANOVA, electrode x laterality interaction, F (1, 28) = 3645.950, p < 0.001). The pattern of polarization was apparent in individual trials (Fig. 2C). An ROC analysis showed that polarization in both electrodes was larger for contralateral saccades, distinguishing this preferred direction from the onset of the preparatory polarization (One-way independent group t-test, F3: t (28) = 22.254, p < 0.001; F4: t (28) = 24.758, p < 0.001).

### Frontolateral ramping polarization varies with response latency

We observed frontolateral ramping polarization that varied with saccade response latency. Polarization aligned to target onset revealed clear differences based on upcoming saccade time: trials with earlier saccades exhibited larger polarizations after the target in both electrode sites (Fig. 3AB, left; Repeated-measures ANOVA, F3: F(1, 28) = 22.236, p < 0.001; F4: F(1, 28) = 56.042, p < 0.001). However, polarization aligned to saccade execution showed mixed results. At F3, slower saccades were associated with higher polarization, whereas no significant differences were observed at F4 (Fig. 3AB, right; Repeated-measures ANOVA, Repeated-measures ANOVA, F3: F(1, 28) = 15.168, p < 0.001; F4: F(1, 28) = 1.191, p = 0.284).

**Fig. 3.**
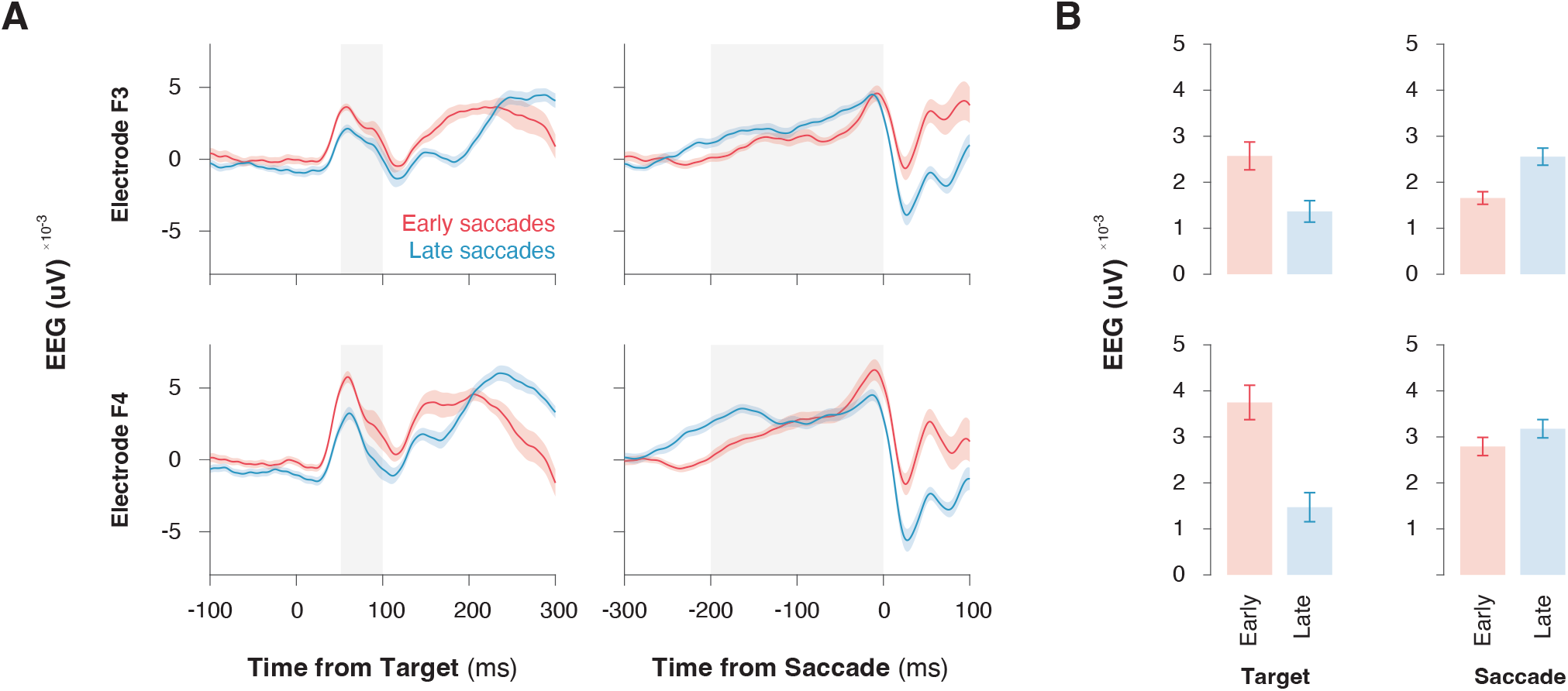
Frontolateral event-related potentials are modulated by response latency. **A**. The lateralized readiness potential observed at electrodes F3 and F4, aligned to target onset (left) and saccade onset (right), for early saccades (red) and late saccades (blue). **B**. Bar plots comparing mean EEG amplitudes for early and late saccades during the target onset and saccade epochs. Error bars represent standard error of the mean (SEM).

### Saccade readiness potential precedes non-goal-directed saccades

To understand the function of this preparatory potential, we considered three alternative hypotheses. First, if these preparatory potentials reflected goal-directed saccade planning processes specifically, then they should only be present for saccades to a cued target and not to other saccades that may have occurred during the session. Alternatively, if these signals are specific to self-initiated movement, then the signal should only be present during saccades that occur with no cues. Finally, if this signal is simply a saccade preparation signal, then it should be evident before both goal-directed and non-goal-directed saccades.

Here, we defined goal-directed saccades as eye movements that are directed toward the target – this target served as a cue to initiate the voluntary movement (Fig. 1C, left), and non-goal-directed saccades were eye movements of approximately the same amplitude and direction that are un-cued and not rewarded (Fig. 1C, right). These saccades were sampled from the inter-trial interval during which the monkey could move gaze freely around the screen. No stimuli or other events occurred during this time. We observed saccades with such parameters in 26 out of 29 sessions.

We observed preparatory polarization in frontolateral electrodes during non-goal directed saccades (Fig. 4A) that mirrored the preparatory polarization observed during goal-directed saccades (Fig. 2A): differential polarization began 300 ms prior to the saccade execution and was lateralized for contralateral saccades (Repeated-measures ANOVA, electrode x laterality interaction, F (1, 25) = 1906.544, p < 0.001). This suggests that the signal we observed in EEG is congruent with the saccade preparation hypothesis and is not sensitive to the motivation of the saccade.

**Fig. 4.**
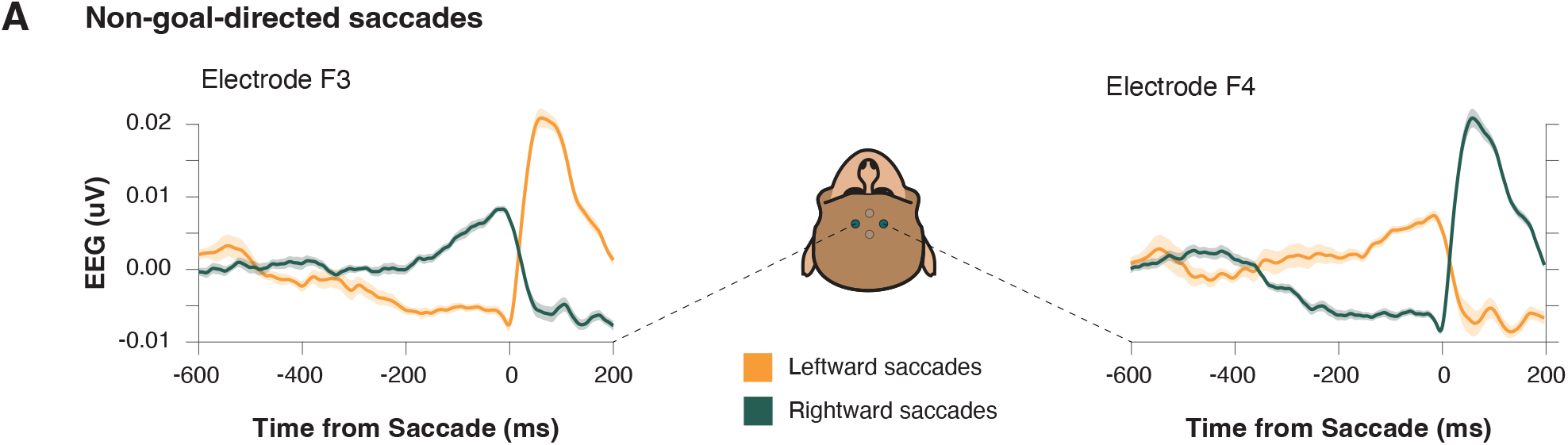
Preparatory activity occurs for non-goal-directed saccades. **A**. Trial-averaged event-related potentials from an example session for left (F3) and right (F4) frontolateral electrodes during leftward (orange) and rightward (green) non-goal-directed saccades. Shaded areas represent standard error of the mean (SEM).

### Event related potentials over frontolateral cortex do not distinguish between stopping and going

So far, we have demonstrated that event-related potentials over the frontolateral cortex ramp until the execution of a saccade, much like FEF neurons (Hanes et al., 1998; Hanes & Schall, 1996). However, earlier work has also demonstrated that FEF also contributes to response inhibition – the cancellation of a planned saccade (Hanes et al., 1998). This earlier work demonstrated that separate populations in the frontal eye field are considered to contribute to the generation and inhibition of a saccadic eye movement.

First, we compared polarization between those trials in which movements were inhibited (canceled) and those in which movements were generated but would’ve been inhibited had a stop-signal appeared (latency-matched no-stop trials). We limited our analyses to trials in the preferred direction for each electrode. Stopping occurs following a stop-signal and is completed within the stop-signal reaction time. Resultantly, if neural signals differentiate between going and stopping should occur within this timeframe. Fig. 5A shows the grand-average ERP for no-stop and canceled trials. Although polarization appeared to differentiate between trial types around 200 ms after the stop-signal, polarization during the STOP period for both F3 and F4 electrodes did not differentiate between trials in which a movement was generated or inhibited (Fig. 5A, right, one-way repeated-measures ANOVA, F3: F (1, 28) = 0.426, p = 0.519; F4: F (1, 28) = 0.540, p = 0.469).

**Fig. 5.**
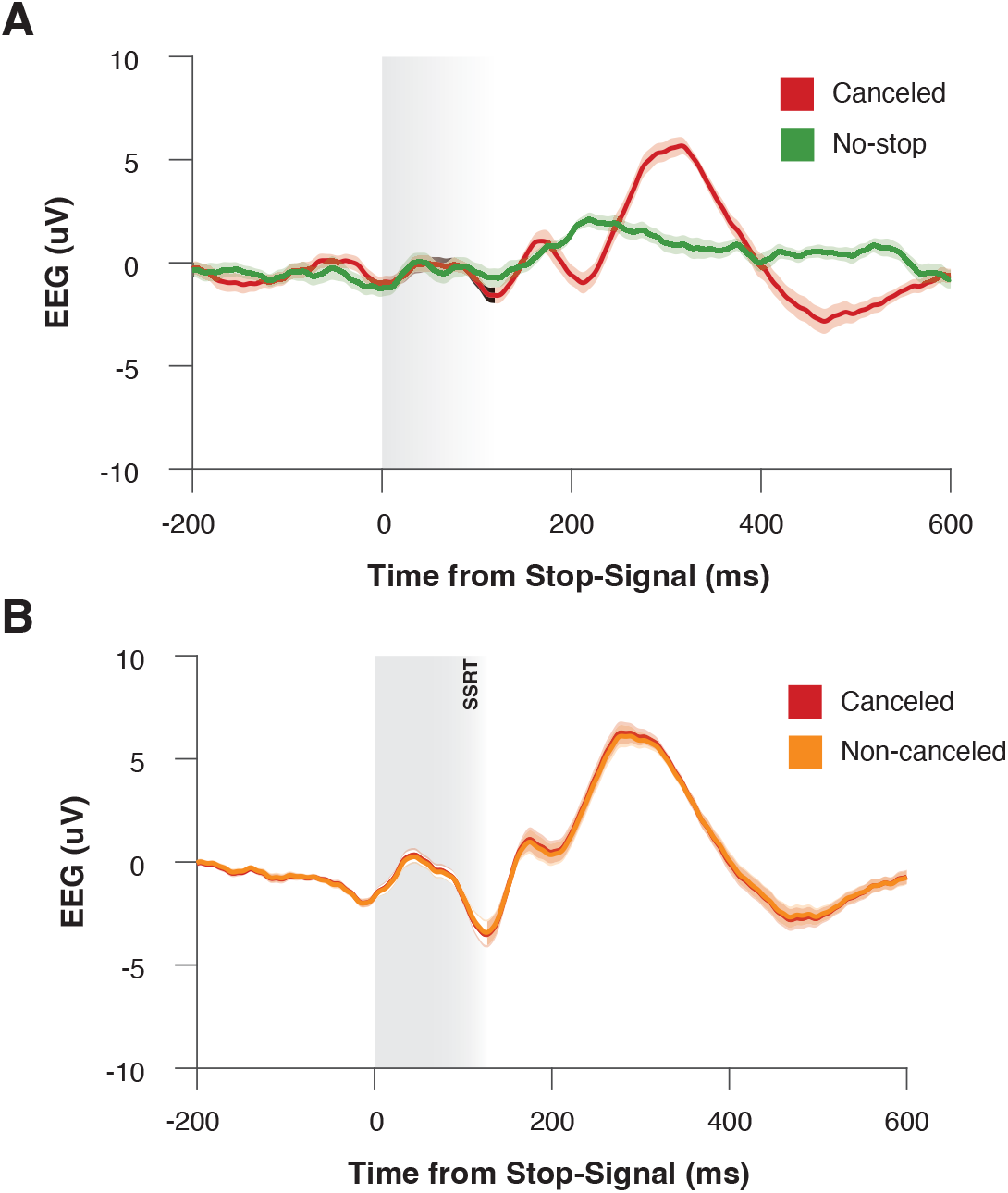
LRP does not index stopping. **A**. Trial-averaged event-related potentials (ERPs) from an example session, during instances of saccade generation (green) and saccade inhibition (red). Data is aligned on the time at which the stop-signal appeared (in canceled trials) or would have appeared (on no-stop-signal trials). Data is collapsed across both electrodes for the preferred saccade direction. Shaded areas represent standard error of the mean (SEM). **B**. Trial-averaged event-related potentials (ERPs) from an example session, during instances in which a saccade was successfully inhibited (red) or incorrectly generated (yellow) after the presentation of a stop-signal. Data is aligned on the time at which the stop-signal appeared and is collapsed across both electrodes for the preferred saccade direction. Shaded areas represent standard error of the mean (SEM).

However, although comparisons between these trial types are valid in the context of the race model, they are confounded by task design. Within our task, the stop-signal was the reillumination of the central fixation spot; as such, signals on canceled trials are contaminated by visual artifact which was not present in no-stop trials. To address this, we compared polarization during the STOP period between non-canceled and canceled trials. Whilst this approach is confounded by RT differences as formulated by the race model (Logan & Cowan, 1984), a visual stop-signal is displayed in both trial types. Fig. 5A shows the grand-average ERP for non-canceled trials and latency-matched no-stop trials. Regardless of this potential confound, we observed no significant difference between these trial types (Fig. 5B, one-way repeated-measures ANOVA, F3: F (1, 28) = 4.858, p = 0.036; F4: F (1, 28) = 0.115, p = 0.737). These findings suggest that frontolateral EEG may not be a useful index of saccade response inhibition.

## DISCUSSION

### Summary of findings

In this study, we investigated saccadic eye movements in monkeys and made several key findings. We observed that monkeys were able to produce planned saccades to targets and could inhibit them when instructed. Monkeys also made non-goal directed saccades during the inter-trial interval, with a portion of these saccades exhibiting characteristics similar to goal-directed saccades. Event-related potentials (ERPs) recorded over the frontolateral cortex showed a gradual ramping polarization prior to saccade execution, consistent with the readiness potential observed in humans. This preparatory potential was present for both goal-directed and non-goal-directed saccades. However, the ERPs we observed could not distinguish between stopping and going, suggesting limited utility for differentiating saccade generation and inhibition.

### Ramping polarization and saccade preparation

Previous work has identified several pre-saccadic potentials in the EEG signal, providing insight into the neural mechanisms underlying saccade preparation. Two primary ramping or preparatory potentials that have been extensively studied are the contingent negative variation (CNV) and the lateralized readiness potential (LRP). The CNV, a slow negative-going waveform, emerges during the interval between a warning signal and a subsequent imperative cue. It signals general anticipatory and preparatory processes leading up to saccade initiation, reflecting broad attentional and cognitive preparation. This potential is observed as a wide distribution of polarization across frontal, central, and parietal regions (Klostermann et al., 1994; Walter et al., 1964). In contrast, the LRP indexes more specific motor preparation, as it captures the buildup of lateralized motor activity over the contralateral cortex before saccade initiation. The LRP is a more localized signal, occurring 200–300 milliseconds before saccade onset, and reflects the selection and readiness of the motor system to execute the movement (Balaban & Weinstein, 1985; Brickett et al., 1984; Evdokimidis et al., 1991; Evdokimidis et al., 1992; Everling, Krappmann, & Flohr, 1997; Everling, Krappmann, Spantekow, et al., 1997; Kurtzberg & Vaughan, 1982; Moster & Goldberg, 1990; Thickbroom & Mastaglia, 1985). Thus, while the CNV is primarily associated with broad attentional preparation, the LRP specifically reflects the motor response preparation and the readiness of the motor system to initiate the saccade. Together, these two potentials represent the ramping of neural activity in anticipation of a forthcoming action—one related to general readiness (CNV) and the other to motor-specific readiness (LRP).

In this study, both monkeys in the study exhibited a clear ramp in polarization at frontolateral electrode sites following an initial visual transient and prior to the execution of a saccade, with the potential showing a clear lateralization—stronger positivity contralateral to the saccade. These observations are similar to those made by Sander et al. (2010), who reported strongest positivity contralateral to the saccade. Interestingly, in contrast to humans, where saccade readiness potentials are typically characterized by a posterior negativity (Everling, Krappmann, & Flohr, 1997; Moster & Goldberg, 1990; Thickbroom & Mastaglia, 1985), the monkey data suggest a distinct frontolateral positivity. This flipped polarity of event-related potentials between humans and macaques has been observed with other signals such as the N2pc (Woodman et al., 2007). Differences in the organization of sulci and gyri, patterns of cortical folding, and the timing of neural signal propagation between humans and macaques are likely sources of the variation in ERP polarity.

Interestingly, the onset of this potential in monkeys at approximately 300 ms before saccade onset aligns with the buildup polarization previously observed in frontal eye field (FEF) neurons during the saccade countermanding task (Hanes et al., 1998). In saccade countermanding tasks, neurons in the FEF exhibit a ramping increase in activity in the period leading up to the saccade, and this increase is thought to reflect the process of accumulating evidence to make a motor decision (i.e., to initiate a saccade). If the decision to move is inhibited, the buildup of activity is halted, and the saccade does not occur. Similarities in the timing between EEG and FEF ramping activity have also been noted by Sander et al. (2010) who describe their positive potential in monkeys beginning to ramp around 100 ms prior to saccade onset; although earlier than those observed here, these latencies were also described as similar to those of FEF neurons during pro- and anti-saccade trials recorded in the same lab (Everling & Munoz, 2000). As such, the potential we have observed in our EEG signal may reflect the synchronized activity of a network of neurons, including those in the FEF, as they ramp up toward threshold for generating a saccade.

After establishing this signal in saccade preparation, we set out to determine if these signals only occur for goal-directed saccade planning, self-initiated movement, or general saccade preparation. Goal-directed saccades are intentional, targeting specific locations, while non-goal-directed saccades occur reflexively or spontaneously. If tied to goal-directed planning, these signals should appear only during saccades to a cued target. If specific to self-initiated movements, they should occur only during uncued saccades. Alternatively, if they represent general saccade preparation, they should be present for both goal-directed and non-goal-directed saccades. Supporting these distinctions, previous work has shown that macaque frontal eye field (FEF) neurons discharge before visually guided and memory-guided saccades (Bruce & Goldberg, 1985), even in darkness, but not before spontaneous saccades in similar conditions (Bizzi, 1968). However, recent work challenged this latter finding demonstrating that FEF neurons do discharge before non-goal-directed saccades, though were more variable in their trial-to-trial firing rate, had unchanged β-band power, and demonstrated shorter timing from direction selection to saccade onset compared to goal directed saccades (Sendhilnathan et al., 2021). Supporting this, we also observed preparatory polarization in the EEG signals for both goal-directed and non-goal-directed saccades, suggesting that this polarization may serve as a general index of saccade preparation.

Although indirectly tested in this study, there are clear parallels between preparatory ramping activity within the frontal eye field, and the polarization observed in frontolateral EEG contacts. Identifying such markers of intracranial activity are important, as these methods are less invasive and do not require any surgical intervention, reducing the risk and discomfort for the participants. This is particularly important when studying human subjects, as invasive measurements may not be feasible or ethically justifiable. Furthermore, non-invasive techniques are typically more widely accessible and can be applied to a larger population, including individuals with certain health conditions or those who cannot undergo invasive procedures. However, future studies should validate such methods by simultaneously recording intracranial and extracranial signals and investigating the biophysical relationship between them.

### Frontolateral event-related potentials do not distinguish between stopping and going in monkeys

Interestingly, although we found preparatory polarization for generating a saccade, we were unable to demonstrate activity that distinguished between stopping and going. When studying motor responses to visual cues, visual-evoked potentials (VEPs) can contaminate the ERP signal, as the onset of the visual cue triggering the motor response also elicits a VEP. This can lead to the contamination of the motor ERP by the VEP, making it difficult to isolate and interpret the neural polarization associated specifically with the motor response. First, although not a valid approach in the context of the race model, if signals from these frontolateral electrodes could differentiate between stopping and going, we may have expected to have seen some distinction between canceled and non-canceled trials. In both instances, a visual stop-signal was presented but in one instance this stop-signal resulted in successfully halting the planned saccade and the other instance resulted in a saccade being executed. Despite these two outcomes sharing the VEP confound, there was no significant difference between the two conditions during the period in which a saccade could be inhibited. Second, there may be biophysical parameters that constrain the contribution of one population of neurons within FEF to the EEG signals recorded over the scalp. Notably, stopping may be achieved by fixation neurons reactivating after the stop-signal is presented, exerting strong inhibition over movement neurons within FEF. The limited contribution of these fixation neurons is two-fold: first, fixation neurons are small and may be inhibitory interneurons – as such, their small size and biophysical properties make it unlikely that they can meaningfully contribute to the EEG signal; second, fixation neurons may be located within the fundus of FEF – as such, even if they were larger pyramidal cells, their dipoles will be orthogonal to movement neurons which are located within the dorsal bank and thus may not be captured as efficiently by EEG.

Interestingly, in human studies of countermanding, several EEG markers have been identified to differentiate between stopping and going and have helped form the basis of neurocognitive models of response inhibition (Diesburg & Wessel, 2021). After sensory processing in the rIFG, a cascade of activity leads to the inhibition of motor plans, with frontocentral low-frequency activity (theta band; ∼2-8Hz) reflecting increased activity in medial frontal areas like the anterior cingulate cortex and pre-SMA during inhibition (Nigbur et al., 2011). The P3, a prominent frontocentral ERP, is particularly notable in studies of stopping (Sajad et al., 2022), with earlier P3 onset latencies correlating with faster stop-signal reaction times (SSRT) and better stopping performance (Kok et al., 2004; Ramautar et al., 2004; Wessel & Aron, 2015). Other studies show a clear relationship between P3 amplitude and motor activity, with larger P3s associated with lower levels of prepotent motor activity and more successful inhibition (Wessel, 2018). Perhaps most relevant, β-bursts over medial frontal and sensorimotor cortices have also been linked to inhibition of motor plans. Wessel (2020)demonstrated that β-bursting decreases steadily over bilateral sensorimotor sites during motor preparation, signaling the inhibition of the motor system that must be overcome to execute the movement. In contrast, during successful movement cancellation, β-bursts increase over frontocentral sites, followed by a short-latency rise in sensorimotor β-bursts, indicating the rapid reinstatement of motor inhibition by frontal areas. In our earlier investigations, we also observed an increase in frontocentral β-bursts during stopping but previously concluded that they are unlikely to contribute to reactive inhibition due to their incidence and lack of specificity (Errington et al., 2020). Given the new findings of sensorimotor readiness potentials in frontolateral sites, future work could re-examine this line of research to determine if β-bursts are more closely linked to distinguishing between stopping and going within these areas instead.

Through this manuscript, we have established compelling evidence for the presence of a preparatory potential in general saccade production, which is a promising finding for understanding the neural mechanisms underlying motor planning. Similar approaches establishing ERPs in the macaque model have been fruitful previously. In our earlier studies we established that monkeys exhibit other ERP’s such as the ERN and an N2/P3 complex over the medial frontal cortex during the saccade countermanding task (Sajad et al., 2022; Sajad et al., 2019). By combining this data with intracranial laminar recordings, we have been able to develop biophysical models of these processes that allow us to understand how single neurons can contribute to larger scale dynamics (Herrera et al., 2023; Herrera et al., 2022). Although such approaches for the medial frontal cortex are more straightforward due to the favorable geometry of the cortex relative to the EEG sensors, we hope that our findings will drive future research to simultaneously record from the FEF whilst capturing EEG to further our understanding of saccade generation and inhibition.

## ACKNOWLEDGEMENTS

This work was supported by NIH R01-EY019882 and P30-EY08126.

## Notes

### Competing Interest Statement

The authors have declared no competing interest.

